# Synaptophysin is a β-Amyloid Target that Regulates Synaptic Plasticity and Seizure Susceptibility in an Alzhiemer’s Model

**DOI:** 10.1101/129551

**Authors:** Daniel J. Adams, Chong Shen, Josien Levenga, Tamara Basta, Stephen P. Eisenberg, James Mapes, Lukas Hampton, Kelly Grounds, Charles A. Hoeffer, Michael H. B. Stowell

## Abstract

Alzheimer’s disease (AD), a condition characterized by cognitive deficits and progressive loss of memory, is causally linked to the short amyloid peptide Aβ42, which disrupts normal neurotransmission^1,2^. Neurotransmitter (NT) release from synaptic vesicles (SV) requires coordinated binding of the conserved core secretory machinery comprised of the soluble NSF attachment protein receptor (vSNARE) synaptobrevin 2 (VAMP2) on the SV and the cognate tSNAREs on the plasma membrane. Synaptophysin (SYP) is the most abundant SV protein^3^ and the major pre-fusion binding partner of VAMP2^4^. A major challenge in understanding the etiology and prevention of AD is determining the proteins directly targeted by Aβ42 and elucidating if these targets mediate disease phenotypes. Here we demonstrate that Aβ42 binds to SYP with picomolar affinity and disrupts the SYP/VAMP2 complex resulting in inhibition of both neurotransmitter release and synaptic plasticity. While functionally redundant paralogs of SYP have masked its critical activity in knockout studies^5,6^, we now demonstrate a profound seizure susceptibility phenotype in SYP knockout mice that is recapitulated in an AD model mouse. Our studies imply a subtle yet critical role for SYP in the synaptic vesicle cycle and the etiology of AD.

## Results

Soluble Aβ42 oligomers are the primary toxic species driving AD^12^, and mammalian synapses exposed to Aβ42 have slower kinetics of evoked synaptic release^13,14^. We hypothesized that this phenomenon is mediated by the major synaptic vesicle protein SYP, as it is reported to form a large multimeric complex with the essential vSNARE VAMP2 and potentially modulates synaptic vesicle fusion^14–16^. To investigate a role for SYP in Aβ42-disruption of SV release, we first developed a standardized method to reliably produce Aβ42 peptide that remained predominantly as monomers, dimers and soluble oligomers (Extended Data Fig. 1). This ensured consistent use of the disease-relevant form, as Aβ42 neurotoxicity and neuromodulatory activities are critically sensitive to its aggregation state^12,17^. Cultured cortical neurons from WT and *Syp*^*-/-*^ mice were then treated with 15 nM Aβ42 or scrambled control peptide, and the evoked release kinetics of the readily releasable pool (RRP) of SVs were quantified using FM dye destaining^18^. We implemented single synapse kinetic analysis in order to determine the unloading kinetics at each of ∼180,000 individual synapses and expose the distribution within each sample (Fig. 1a, b). As predicted by previous studies^5,19^, the kinetics of NT release were similar in WT and *Syp*^*-/-*^ neurons (Fig.1c blue and black curves) where synapses most frequently unloaded >60% of labeled SV within 45s. However, the presence of Aβ42 drastically reduced the overall release kinetics of WT synapses, roughly doubling the mode of the population to 85s (Fig. 1a). Additionally, the distribution of kinetics at WT synapses treated with Aβ42 was significantly broader (kurtosis=0.49) than in control-treated WT synapses (kurtosis=4.96), indicating that effects of the peptide at this physiologic concentration are heterogeneous across synapses. Furthermore, while a small number of these synapses retained normal fast kinetics, we also observed a population of synapses with very slow kinetics (τ>400s) in the Aβ42 treatment group in which exocytosis appeared completely halted, as the decay rates were similar to that of background photobleaching (Extended Data Fig. 2). Remarkably, this dramatic shift in kinetics was completely absent in *Syp*^*-/-*^ neurons treated with Aβ42 (Fig. 1b), indicating that SYP is necessary for Aβ42-induced impairment of vesicle release.

**Figure 1.**
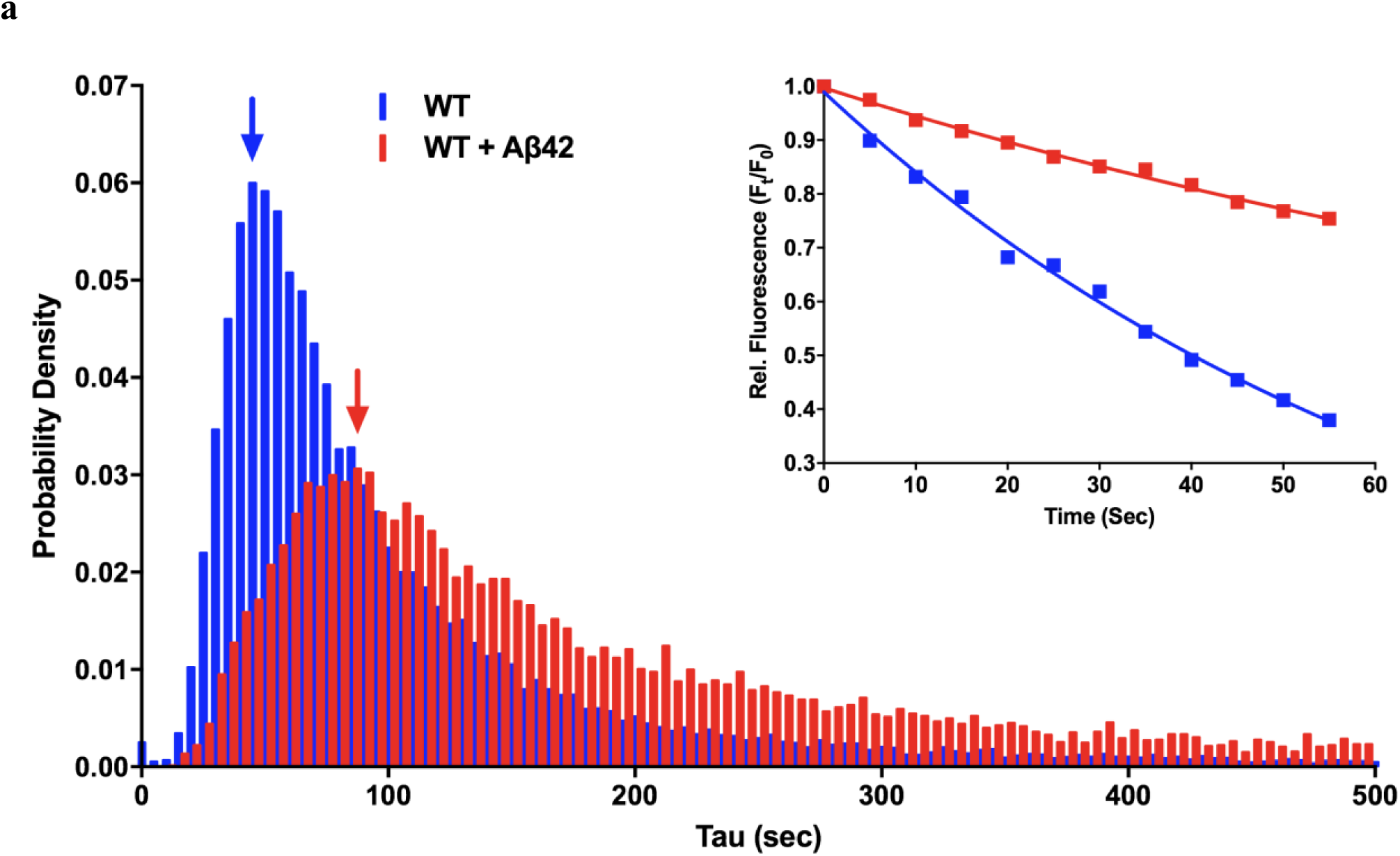
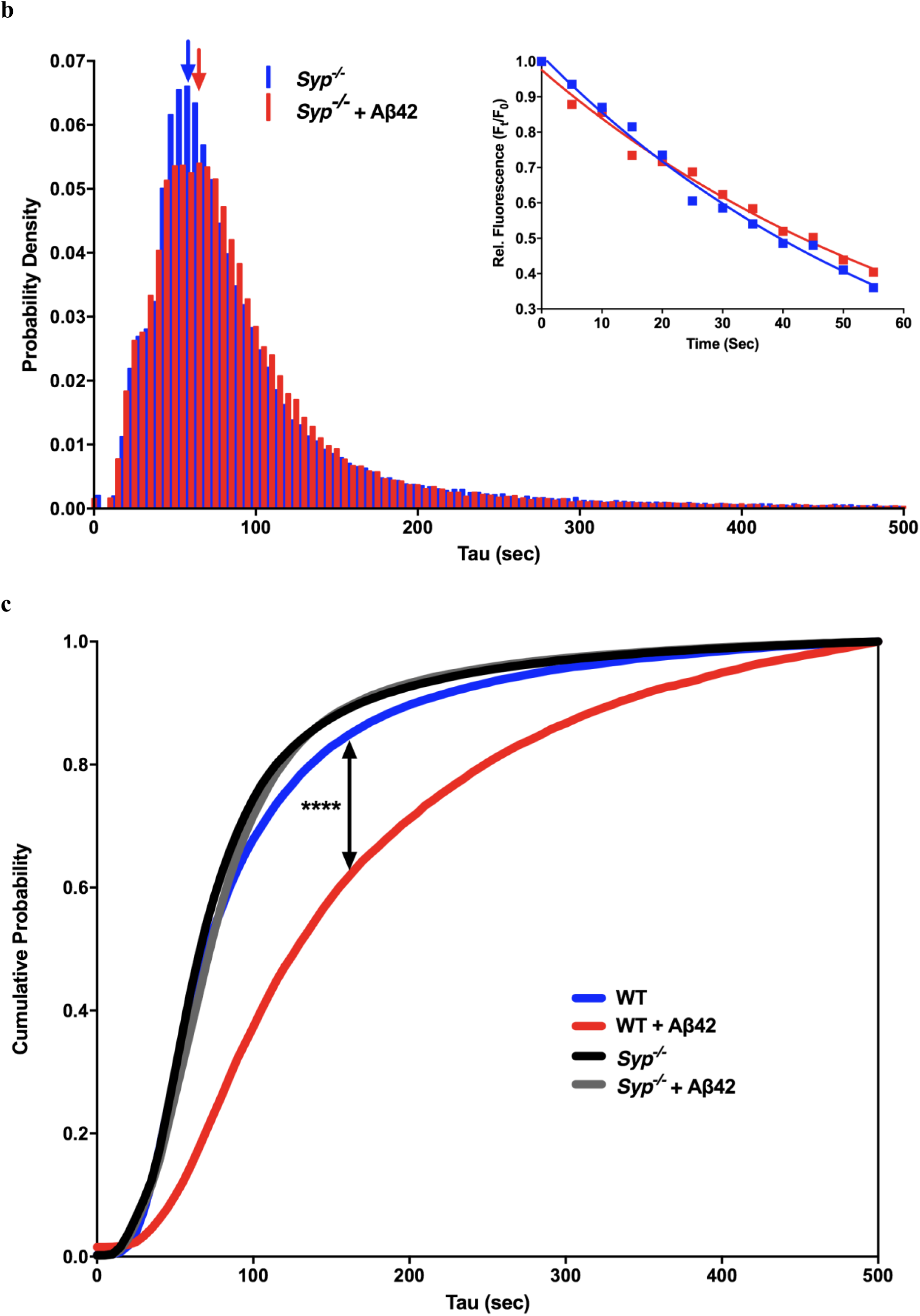
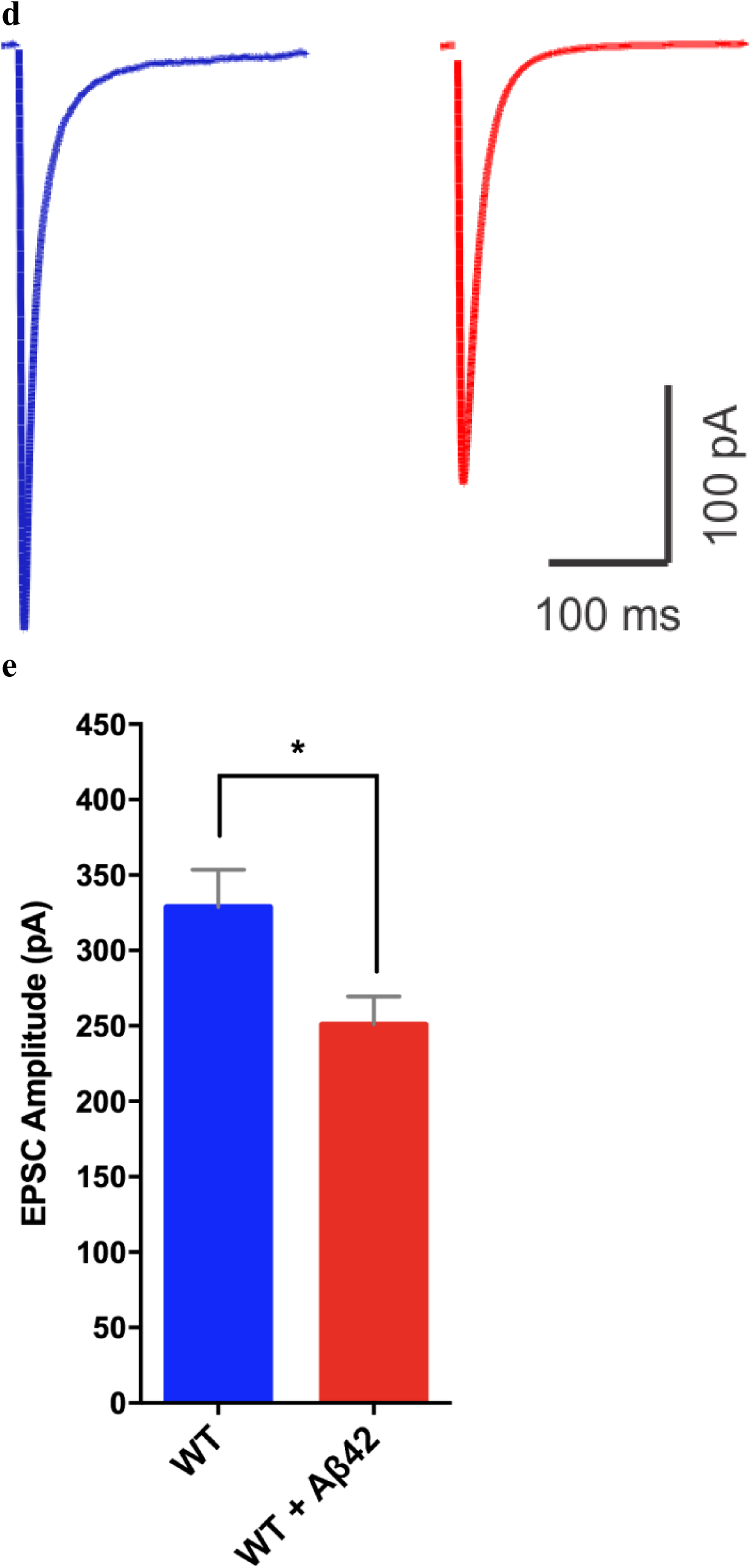
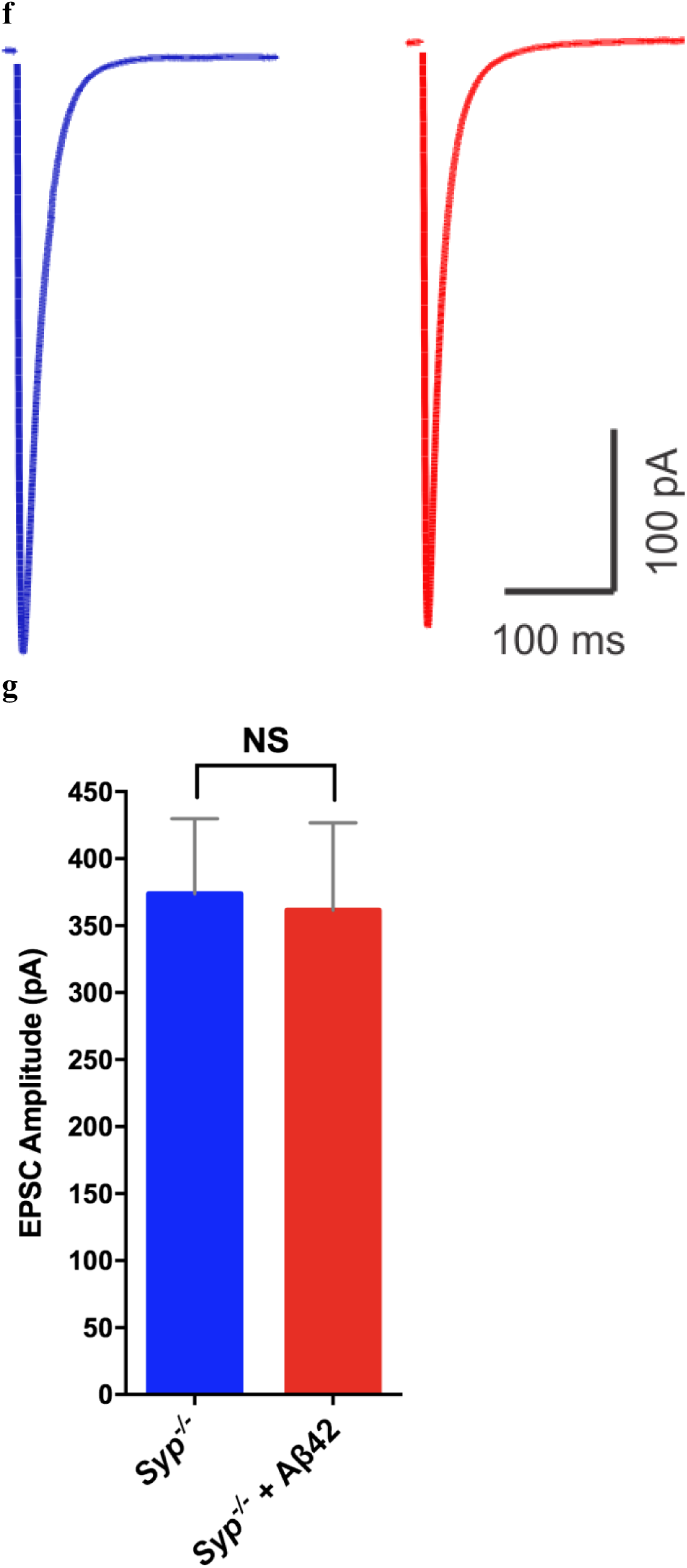
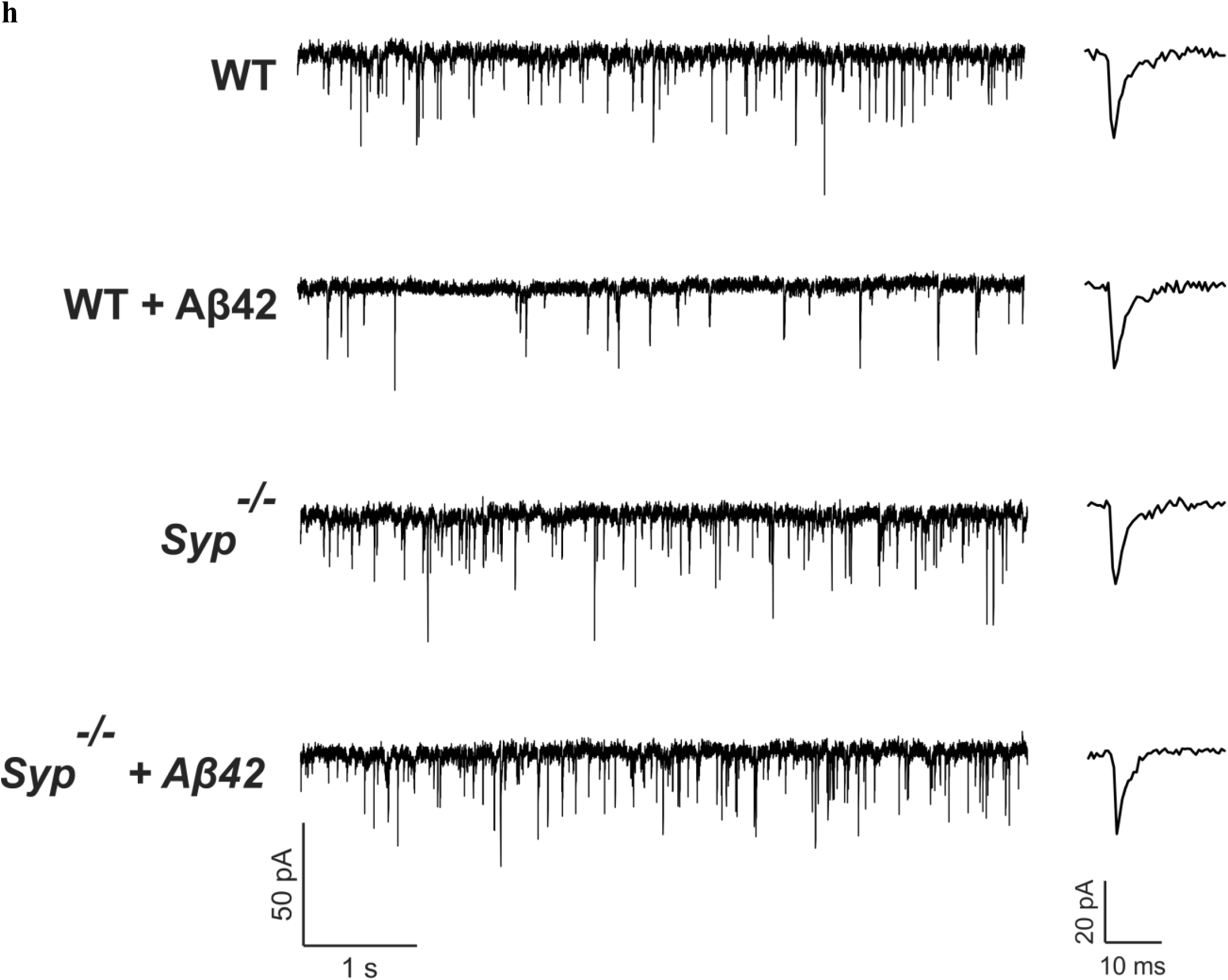
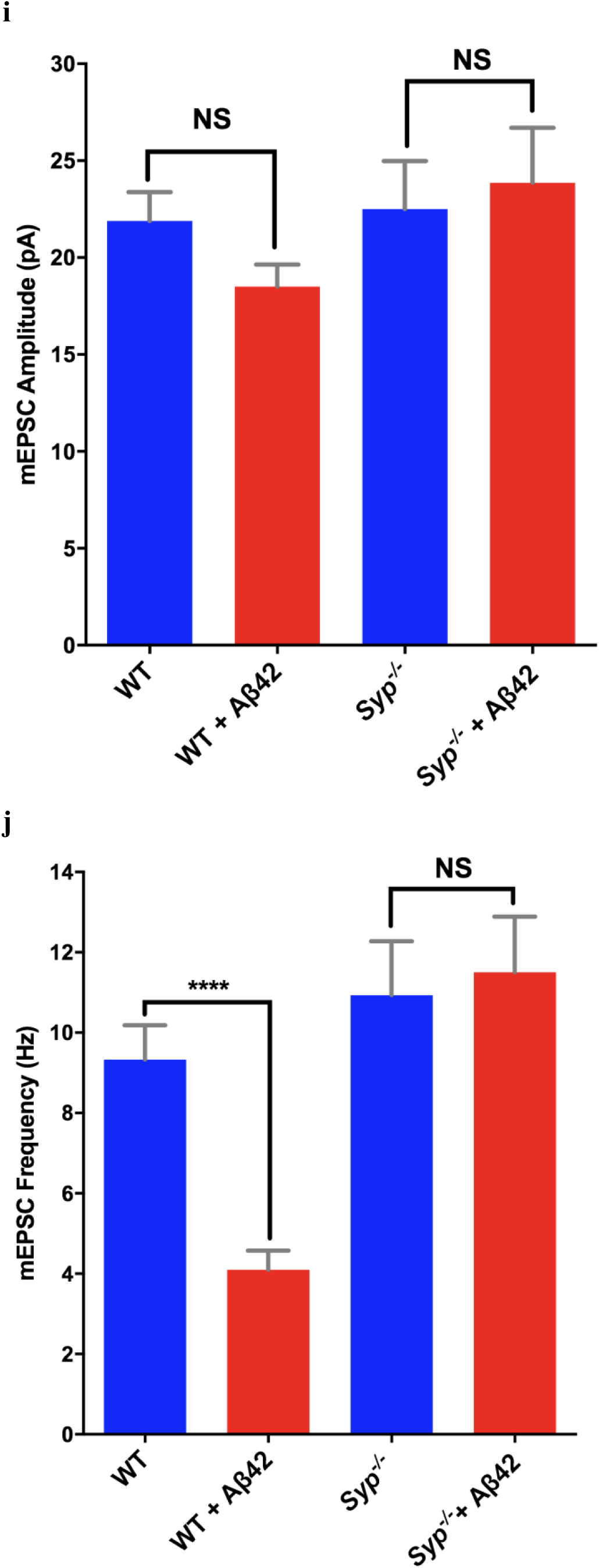
| Aβ42 inhibition of spontaneous and evoked transmission is SYP dependent. **a,b,** Distribution of all kinetic time constants (τ) of FM dye unloading from WT (**a**) or *Syp*^*-/-*^ (**b**) neurons treated with 15 nM Aβ42 (red) or scrambled peptide (blue). Insets show unloading curve from a representative single synapse with fit from bin indicated with arrow (WT + Aβ42, *n*=12,364 synapses from 16 experiments; WT, *n*=22,996 synapses from 15 experiments; *Syp*^*-/-*^ + Aβ42, *n*=68,611 synapses from 14 experiments; *Syp*^*-/-*^, *n*=75,274 synapses from 11 experiments). **c**, Cumulative distribution functions of τ for all 4 conditions. No significant differences between WT, *Syp*^*-/-*^, or *Syp*^*-/-*^ + Aβ42. **d, e,** Averaged evoked EPSC traces (**d**) and average amplitudes (**e**) from WT neurons (blue, *n*=36) and WT neurons treated with 500nM Aβ42 (red, *n*=38). **f, g,** Averaged evoked EPSC traces (**f**) and average amplitudes (**g**) from *Syp*^*-/-*^ neurons (blue, *n*=17) and *Syp*^*-/-*^ neurons treated with 500nM Aβ42 (red, *n*=13). **h**, Representative mEPSC traces from primary WT and *Syp*^*-/-*^ neurons treated ± 500nM Aβ42 with representative spike waveform. **i, j,** Average mEPSC amplitudes (**i**) and frequencies (**j**) of neurons from WT (*n*=19), WT treated with 500nM Aβ42 (*n*=28), *Syp*^*-/-*^ (*n*=18) and *Syp*^*-/-*^ treated with 500 nM Aβ42 (*n*=19). *\*\*\*\*P*<0.0001 (Kolmogorov-Smirnov test) (**c**), *P< 0.01, ****P<0.0001 (unpaired t-test) (**e, g, i, j**). Data are mean±s.e.m.

While FM-dye methodology allowed simultaneous interrogation of a large number of synapses under limited Aβ42 dosing and observation of a comprehensive spectrum of SV release derangements, this approach employed non-physiologic exhaustive stimulation to achieve complete RRP release. To substantiate our findings with a more endogenous form of stimulation, we measured evoked excitatory postsynaptic currents (eEPSCs) in cultured neurons incubated with Aβ42 or scrambled peptide. To ensure that patched cells received adequate treatment, we saturated the culture with the commonly used dose of 500 nM peptide. Treatment with Aβ42 significantly reduced WT eEPSC amplitude, but had little effect on *Syp*^*-/-*^ neurons (Fig. 1d-g). To determine whether the attenuated evoked response arose from reduced release probability, we also measured miniature excitatory postsynaptic potentials (mEPSP) under the similar conditions (Fig. 1h-j). While Aβ42 treatment did not alter spontaneous release amplitude in WT or *Syp*^*-/-*^ neurons, we observed a 56% decrease in mEPSP frequency in WT but not *Syp*^*-/-*^ cells, demonstrating that Aβ42 reduces probability of evoked and spontaneous SV release by a SYP-dependent mechanism.

To explore the possibility that Aβ42 disrupts SV release via direct SYP-Aβ42 binding, we performed surface plasmon resonance (SPR) with Aβ42 or control peptides immobilized as the ligand and assayed binding of purified recombinant human SYP protein as analyte. Using antibodies that selectively bind different Aβ42 aggregation states^20^, we determined that the immobilized peptide was primarily the disease-relevant monomers, dimers and soluble oligomers rather than higher order aggregates (Extended Data fig. 3)^12^. We observed a remarkably high affinity interaction with an apparent K_d_=750 pM for the native SYP hexamer and no observable binding to immobilized control reverse (42-1) peptide (Fig. 2a). As both species were highly purified, this result indicates that the SYP-Aβ42 interaction is direct and can occur in the absence of other proteins or cofactors. Given that the estimated concentration of soluble Aβ42 in brain tissue of AD patients is in the low nanomolar range^21^, we suggest that Aβ42 may bind to SYP in neurons of AD patients to reduce release probability at physiological concentrations.

**Figure 2.**
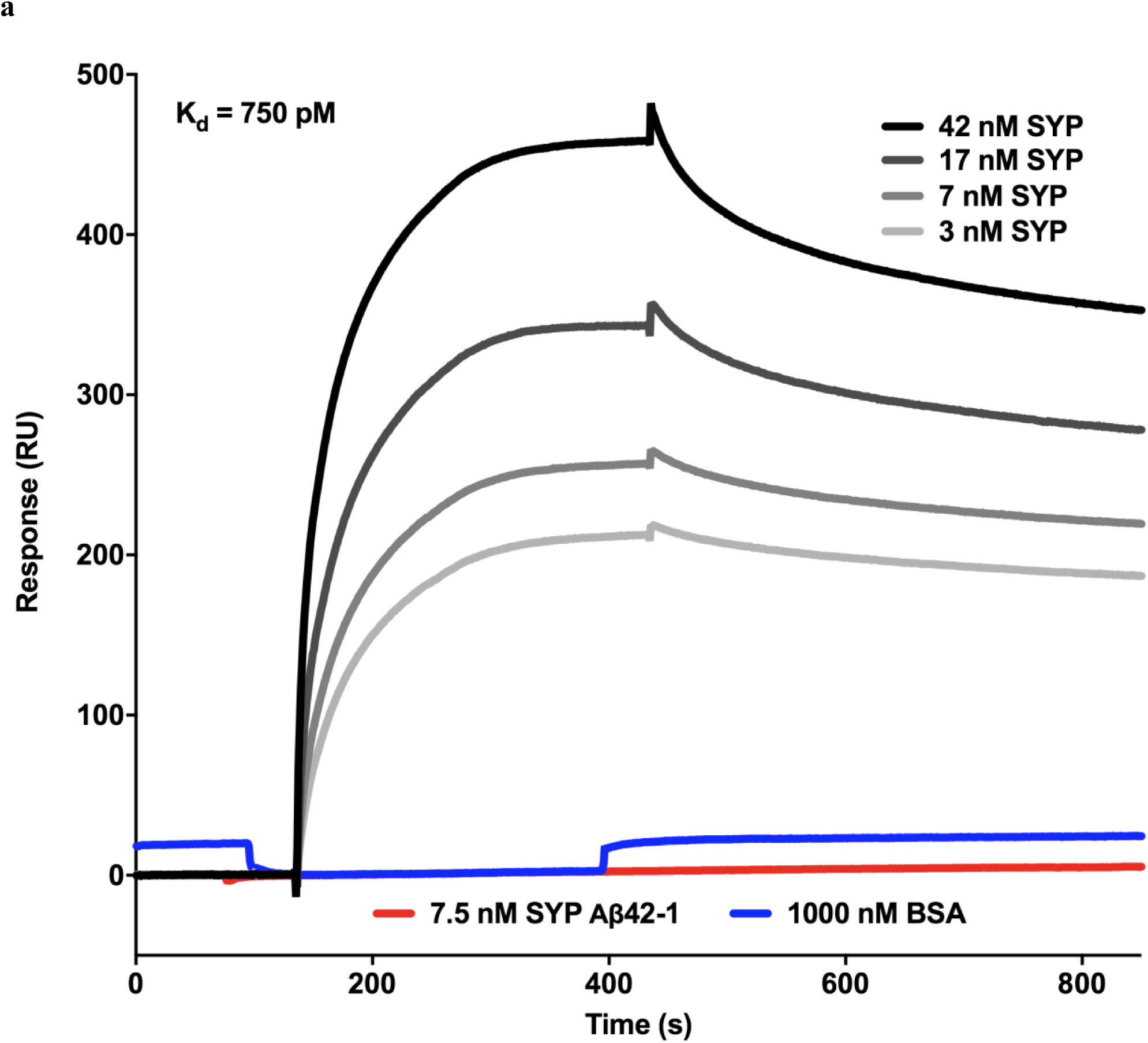
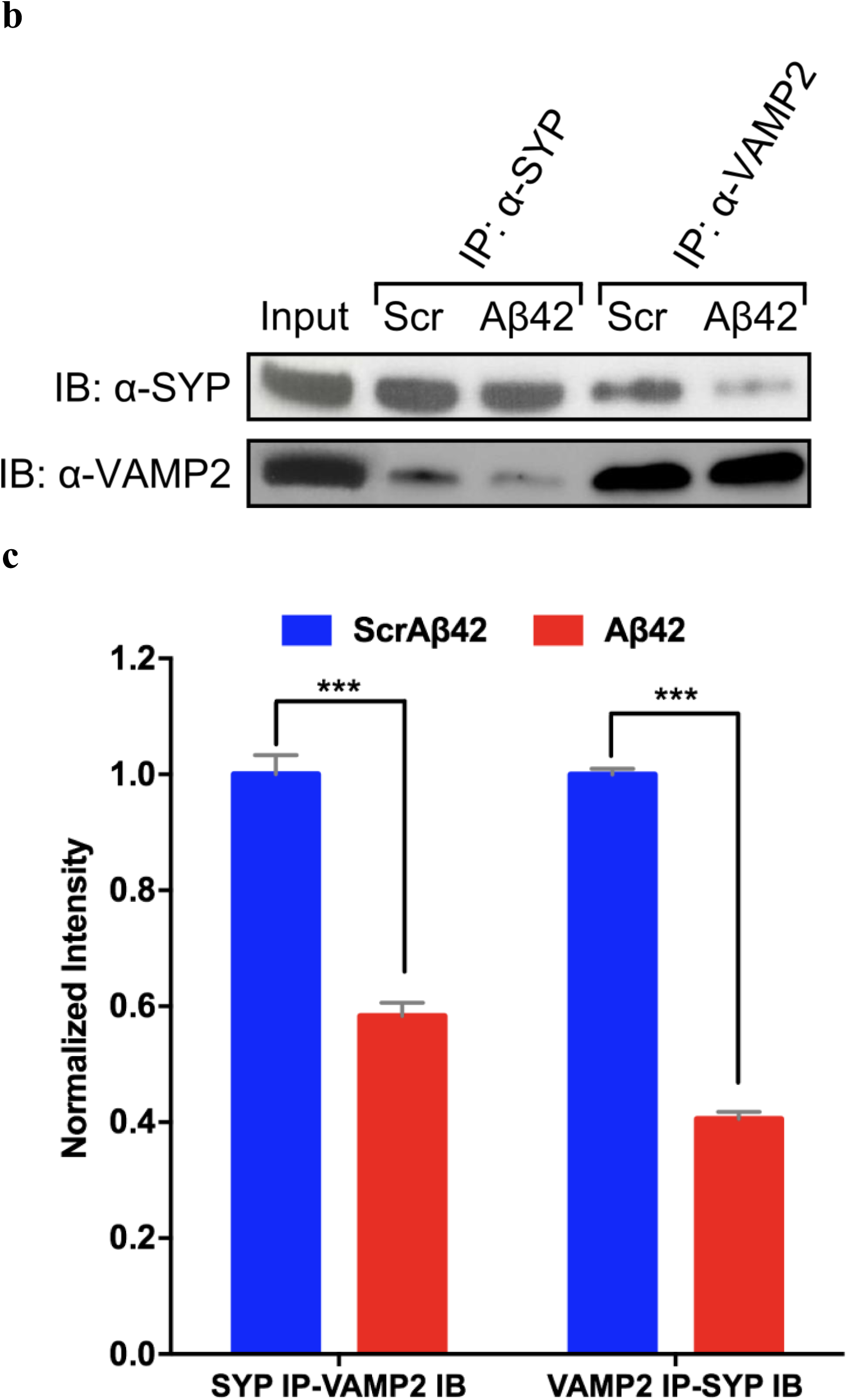
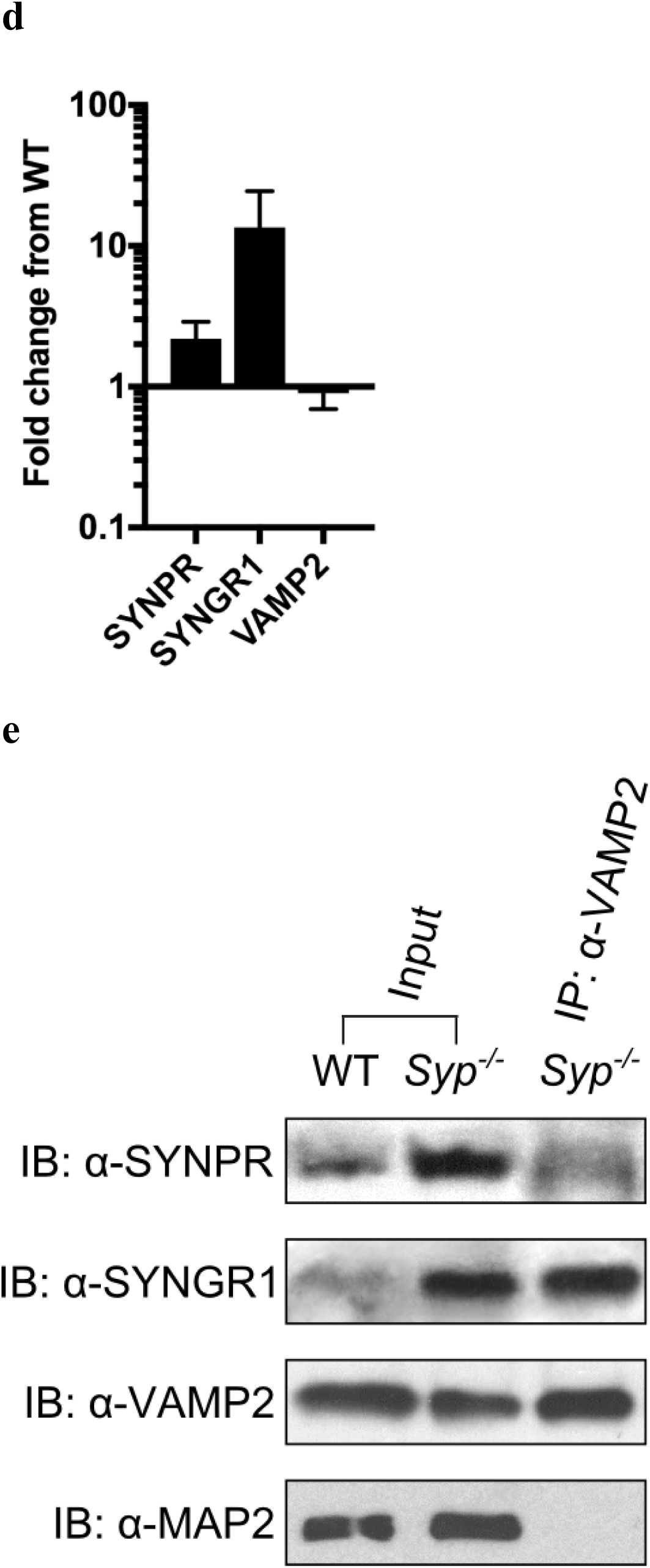
| Aβ42 disrupts the SYP/VAMP2 complex via high affinity interaction. **a,** SPR sensorgrams for binding of recombinant human SYP at several concentrations and BSA to Aβ42; SYP at 7nM binding to reverse Aβ42 (Aβ42-1) in gray. On-rate = 835,507 M-1 sec-1, off-rate = 0.003746 sec-1, calculated K_d_ for the SYP hexamer = 750 picomolar. **b, c,** Cortical neurons treated with 15 nM Aβ42 or scrambled peptide for 24 hrs. were lysed and immunoprecipitated for SYP or VAMP2 and probed for the other protein. Representative blot (**b**) and quantification from multiple experiments (**c**) (SYP IP *n*=3, VAMP IP *n*=2). **d,** Protein levels of SYP paralogs SYNPR and SYNGR1 in *Syp*^*-/-*^ synaptosomal extracts were quantified by immunoblotting (*n=2*) normalized to MAP2 and shown as fold change from protein levels in WT. **e,** *Syp*^*-/-*^ whole brain synaptosomes were immunoprecipitated for VAMP2 and immunoblotted for SYNPR and SYNGR1. *Syp*^*-/-*^ input and WT synaptosomes shown for comparison. **f,** Immunoblot analysis of synaptic proteins bound to column-immobilized Aβ42 or scrambled peptide. MAP2 is used as a loading control. \*\*\**P*<0.001 (t test) (**c**). Data are mean±s.e.m.

Structural features of the SYP/VAMP2 complex suggest that the SYP hexamer enhances the rate of SV fusion by clustering VAMP2 dimers on the SV surface, conferring cooperativity of trans SNARE interactions^15,22^. We posited that a high affinity Aβ42-SYP interaction could disrupt this scaffold function by interrupting SYP/VAMP2 binding. In support of this model, we found that Aβ42 treatment of cultured neurons reduced intact SYP/VAMP2 complexes by ∼50% (Fig. 2b, c). The VAMP2 trans-membrane domain (TMD) is necessary for SYP binding^4^ and manipulations of this TMD also impair vesicle fusion in many model systems^23–25^, consistent with our data suggesting that association with SYP is required for normal release kinetics. Paradoxically, loss of SYP clustering in *Syp*^*-/-*^ neurons appears to have no effect on SV exocytosis^26,27^ (Fig. 1b, f-j), and we hypothesized that related members of the physin protein family may functionally compensate for this loss. The six known neuronal paralogs of SYP feature unusually high conservation across the TMDs (Extended Data Fig. 4a, b) such that these related proteins may bind VAMP2 in the SV membrane and perform the clustering function when SYP is not present. Both synaptoporin (SYNPR) and synaptogyrin 1 (SYNGR1) can compensate for loss of SYP as shown using double knockout mice^5,6^, and we found that both paralogs were significantly upregulated in *Syp*^*-/-*^ brains (Fig. 2d) consistent with the need to replace the ∼30 copies of SYP in each SV^3^. Furthermore, we found that both proteins were bound to VAMP2 in *Syp*^*-/-*^ animals by co-immunoprecipitation (Fig. 2e). However, the lack of sensitivity to Aβ42 observed in *Syp*^*-/-*^ neurons suggests that Aβ42 does not target the SYP paralogs and disrupt binding to VAMP2 as it does with SYP. In support of this, Aβ42 affinity chromatography showed high selectivity of the peptide for SYP over SYNPR or SYNGR1 (Fig. 2f). Additionally, VAMP2 did not co-purify with SYP on the Aβ42 beads suggesting that Aβ42 and VAMP2 competitively bind SYP and that Aβ42 can displace bound VAMP2 via its high affinity interaction with SYP to disrupt its function in vesicle release. In addition to the essential SNARE proteins, a suite of accessory proteins are critically important in activating VAMP2-mediated fusion^7^, and these data suggest that SYP also contributes substantially to this process in a similar role.

Acute Aβ42 treatment of hippocampal slices causes synaptic depression and impairment of LTP, and administration of peptide directly to the brain by microinjection generates short term memory deficits in rats^12^. To determine if such effects on plasticity could also be SYP dependent, field excitatory postsynaptic potentials (fEPSPs) were measured in hippocampal slices from WT and *Syp*^*-/-*^ mice in the presence and absence of Aβ42. Basal synaptic efficacy in *Syp*^*-/-*^ slices was indistinguishable from WT (Extended Data Fig. 5), indicating that there is not a baseline difference in synaptic release probability. However, we found that while Aβ42 strongly inhibited LTP in WT slices, LTP was not significantly affected by Aβ42 in *Syp*^*-/-*^ slices (Fig. 3a-d). This implies that the release defects attributed to loss of SYP/VAMP2 function in cultured neurons (Fig. 1) generate clinically relevant changes in circuit level physiology as well. Failure to form LTP is thought to underlie the cognitive deficits that are hallmarks of early AD^29,30^, implicating loss of SYP activity as an important early event in the disease progression and suggesting that SYP/VAMP2 may be a novel therapeutic target in this unsolved clinical challenge.

**Figure 3.**
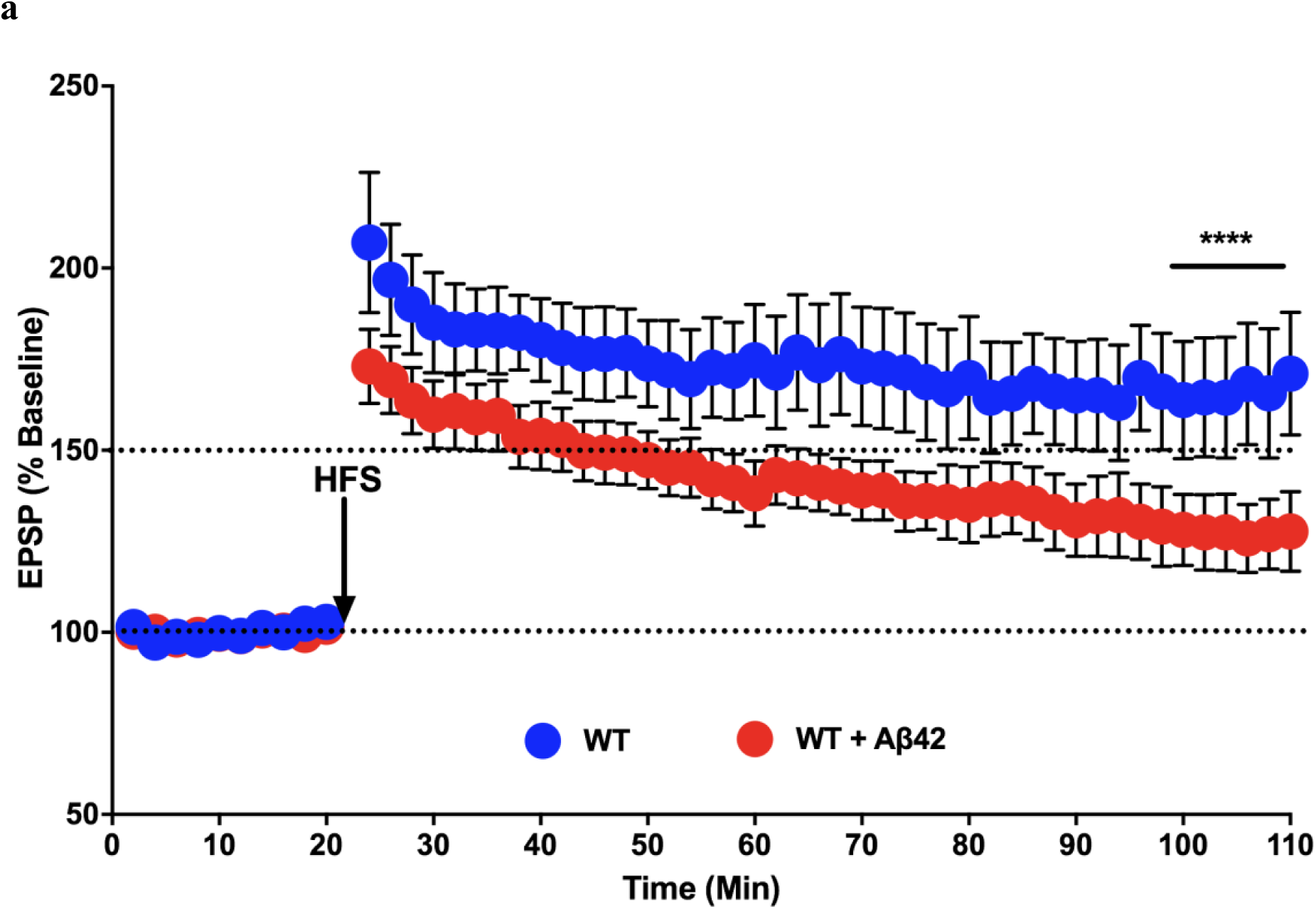
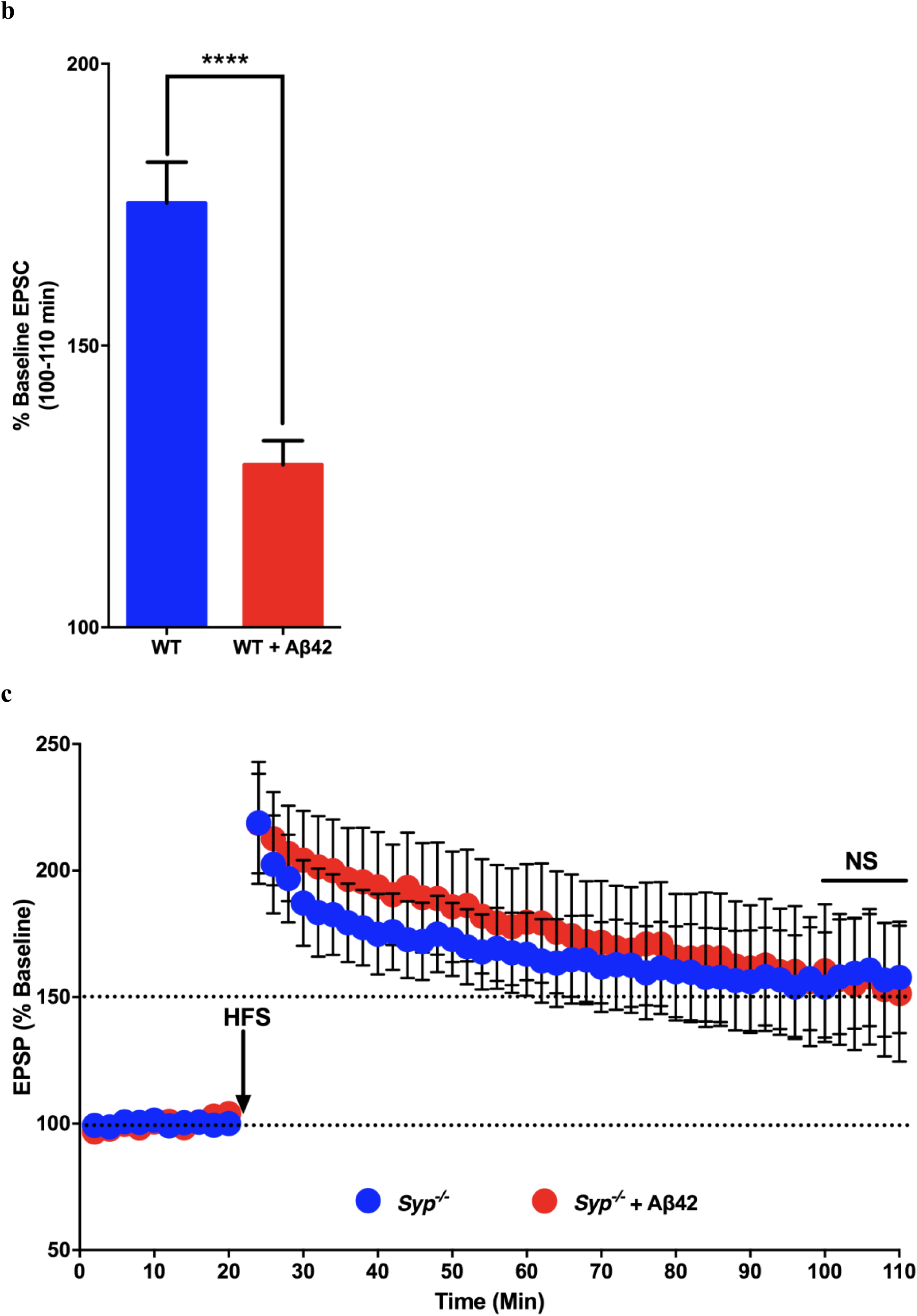
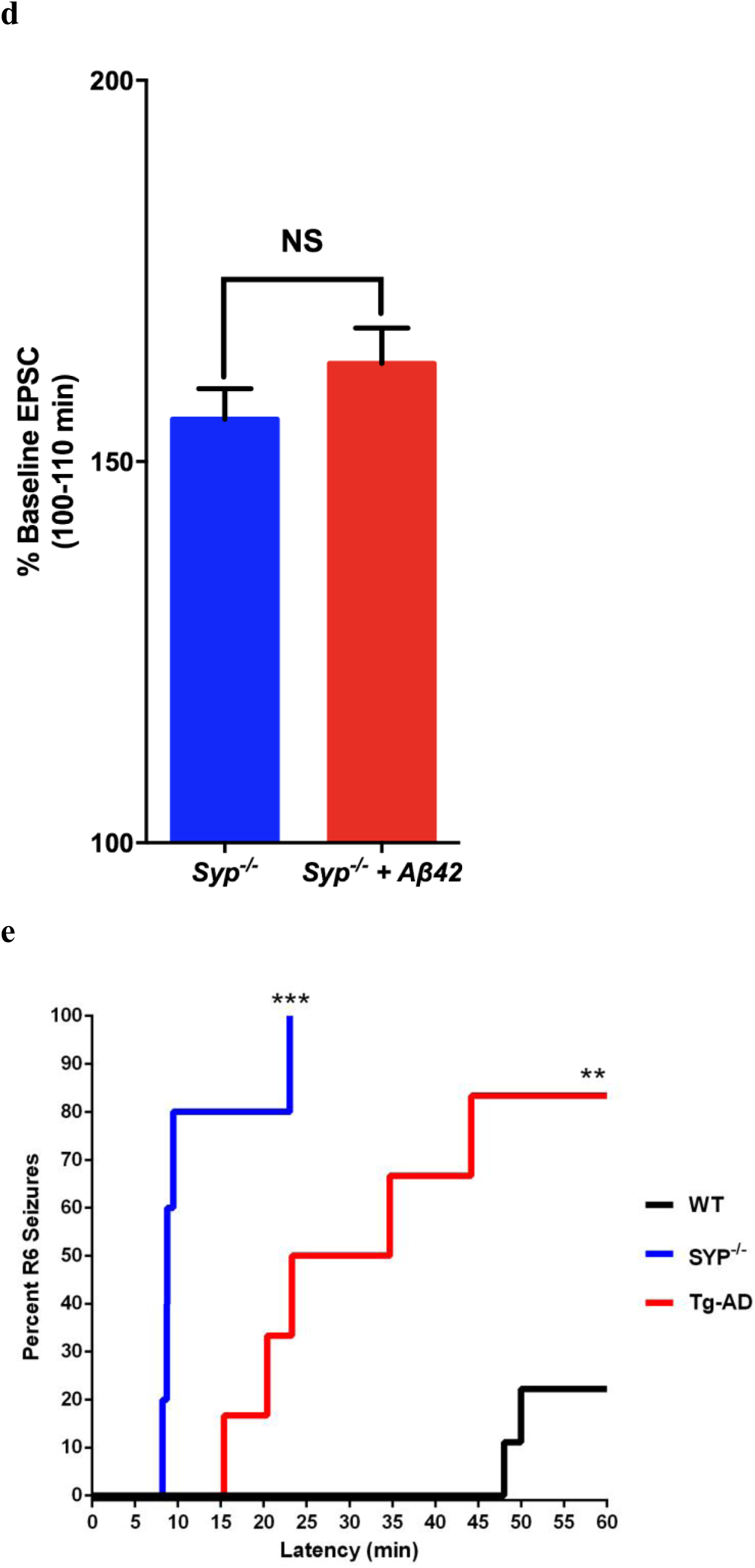
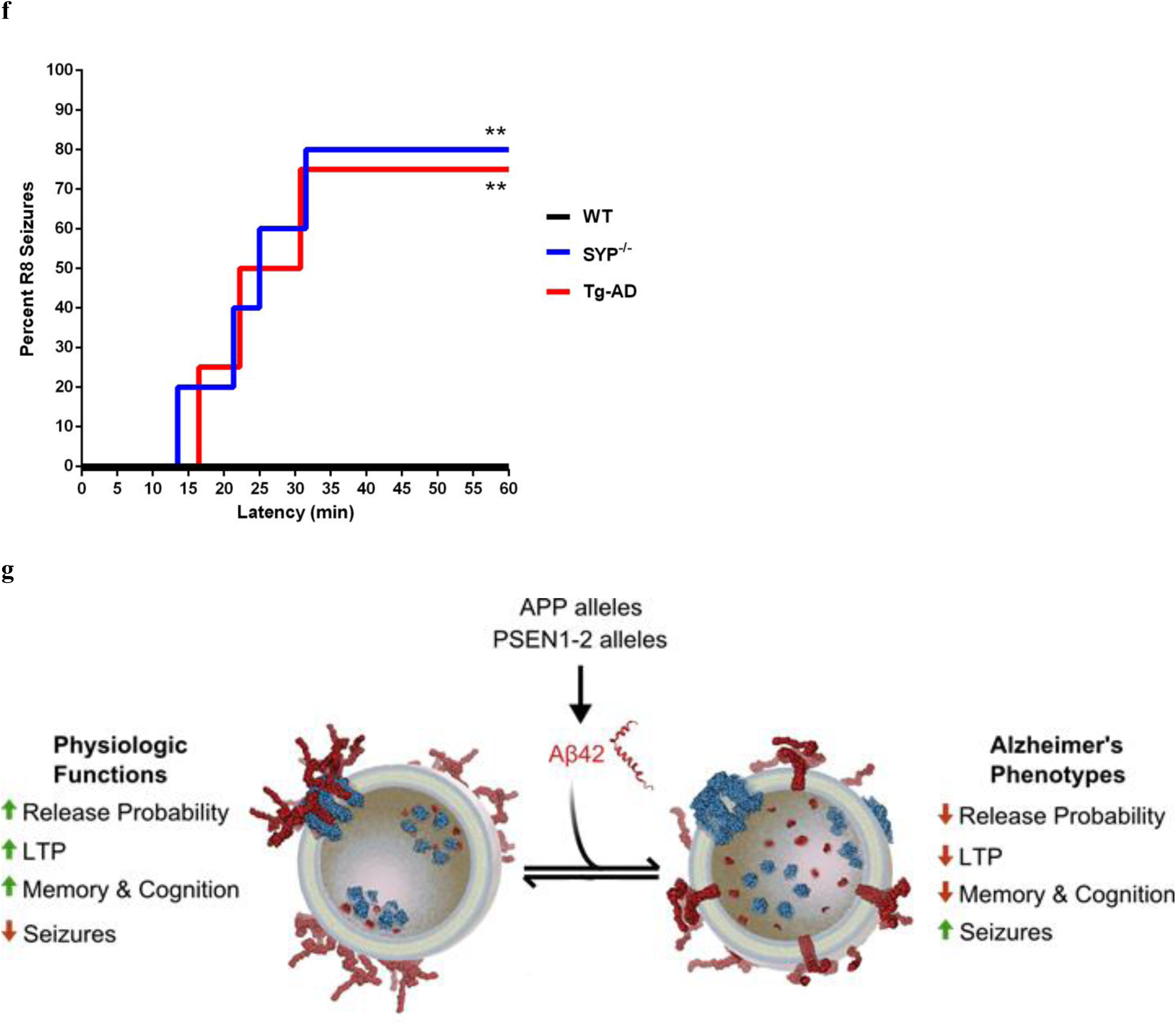
| Loss of SYP/VAMP2 links AD seizures, Aβ42 inhibition of LTP and Alzheimer’s genotypes. **a, b, c, d,** Following 4 hour treatment of hippocampal slices with Aβ42 or control, EPSPs were measured before and after high frequency stimulation. LTP induction in WT (**a, b**, blue, *n*=16) was blocked by treatment with Aβ42 (**a, b,** red, n=16) whereas normal LTP in *Syp*^*-/-*^ (**c, d,** blue, *n*=13) was unperturbed in *Syp*^*-/-*^ slices treated with Aβ42 (**c, d,** red, *n*=11). **e, f,** Mice 4-6 months old of 4 genotypes (WT (*n*=9), *Syp*^*-/-*^ (*n*=5), TgAD (*n*=6), *ApoE4* (*n*=9) were injected with kainic acid (25mg/Kg i.p.) and observed for 60 minutes with video recording. Seizure severity was scored blind by two observers using a modified Racine scale. Latency survival to R6 (**e**) or R8 (**f**) seizure severity are reported. **g,** The strongest AD-associated genotypes lead to increased Aβ42 (*APP* and *PSEN1-2* alleles) which disrupts the prefusion vSNARE complex of SYP (blue) and VAMP2 (homodimer in red) resulting in redistribution of VAMP2 on the SV membrane, subsequent reduction of release probability and dysregulated plasticity. These synaptic defects may underlie disease phenotypes such as seizures and loss of memory and cognition making the SYP/VAMP2 complex a key early target in the disease progression. ****p<0.0001 (unpaired t test with Welch’s correction) (**a, b, c, d**). *\**p<0.05, *\*\**p<0.005, *\*\*\**p<0.0005 (Gehan-Breslow-Wilcoxon test) (**e, f**). Data are mean±s.e.m.

In addition to cognitive and memory phenotypes in AD, co-morbidity of seizures is well documented^31–33^, and we hypothesized that aberrant synaptic release resultant from SYP/VAMP2 disruption may contribute to this pathology. Transgenic AD (Tg-AD) model mice overproducing Aβ42 (Extended Data Fig. 6) recapitulated this clinical feature of the disease, displaying markedly increased susceptibility to kainic acid-induced seizures (Fig. 3e, f). We found that *Syp*^*-/-*^ mice were also highly sensitized to seizures with nearly all of *Syp*^*-/-*^ and Tg-AD mice experiencing clonus followed by death, while no WT animals experienced such seizure severity. Importantly, this is the first observation of a dramatic phenotype in the *Syp*^-/-^ mice and substantiates the role of SYP in synaptic function, at least minimally at the circuit level.

Together, our results inform a model in which Aβ42 directly binds SYP in the SV membrane, displacing VAMP2 dimers, thereby compromising the catalytic function of pre-fusion v-SNARE clustering. When SYP is absent, SYP paralogs are upregulated and bind VAMP2 in its stead to perform this important role in vesicle fusion. Aβ42 does not bind these paralogs and therefore cannot perturb release probability or LTP on the *Syp*^*-/-*^ background. The newly discovered seizure susceptibility phenotype we observe for *Syp*^-/-^ mice suggests at minimus that loss of SYP and subsequent synaptic and circuit level perturbations are an early event in AD.

## Methods

**Animals:** All animal procedures were carried out in accordance with protocols approved by the IACUC at CU Boulder and the Animal Welfare Assurance filed with OLAW. C57BL/6J (B6) mice (Jackson Labs) were used as WT in all experiments. *Syp*^*-/-*^ animals^34^ were a gift of R. Leube at RWTH Aachen University. The TgAD mice (B6C3-Tg(APPswe,PSEN1De9)) were obtained from Jackson Labs.

**Cell Culture:** Cortical neurons were prepared as described previously^35^ and plated at high density (∼5000 cells/mm^2^) to ensure physiologically relevant synaptic connections.

**Aβ42 peptide preparation**: Amyloid peptides were prepared by dissolving a lyophilized film of the peptide at 1mg/ml in 10mM NaOH, followed by bath sonication for 5 minutes and centrifugation at 13,000g for 5 minutes. The concentration was then determined using a Nanodrop at 280 nm and cross-validated using both a BCA assay and SDS-PAGE. Aβ42 samples were further analyzed by native SDS-PAGE and negative stain EM to characterize the oligomeric nature of the amyloid prior to use. All samples were analyzed or utilized within 1 hour of preparation.

**FM 1-43 Assays:** Imaging was performed on primary cortical neurons prepared as above at 12 - 15 DIV. Neurons were treated with 10 to 15 nM Aβ42 or scrambled peptide (R Peptide, Bogart, Georgia or AmideBio, Boulder, Colorado) 24 hours prior to imaging. Cells were treated within 1 hour of sample peptide preparation by removal of 1ml of conditioned media, addition of peptide to the conditioned media and then replacement of the mixture to the culture dish. Neurons were labeled with 10 μM FM 1-43 (Invitrogen, Carlsbad, California) in stimulating buffer (25 mM HEPES pH 7.4, 59 mM NaCl, 70 mM KCl, 2 mM CaCl_2_, 1 mM MgCl_2_, 30 mM glucose) for 2 minutes at 37 °C followed by washing in rest buffer (25 mM HEPES pH 7.4, 124 mM NaCl, 5 mM KCl, 0.2 mM CaCl_2_, 5 mM MgCl_2_, 30 mM glucose) to prevent release of labeled vesicles prior to assay. Cultures were depolarized under profusion with stimulating buffer and imaged for 60 seconds after onset of release. For all experiments synaptic puncta were identified in ImageJ^36^ by making a max projection of the video, background subtraction of 0.5*mean pixel intensity and finding local maxima on a 10 px (∼650 nm) radius with the NEMO-derived^37^ ImageJ plugin 3D Fast Filter. These maps were enlarged over a 5 px radius and mean grey value of each punctum was plotted against time. Each unloading curve was fitted to an exponential decay equation of the form *f_(t)_ = f_0_ · e^(-1/τ)t^ + c* to determine a time constant, τ, to represent the kinetics of release at each synapse. These data were filtered for particles whose behavior poorly fit the exponential model (R^2^<0.95) and the data selected for values of between 0 and 500 seconds, although a small number of extremely slow decay events >500 seconds, were observed. The included τ values were sorted into 5 second bins and displayed as a histogram. Each bin represents the aggregate probability density from all biological replicates at each τ value.

**Single cell recordings:** Cultured cortical neurons prepared as above. Evoked synaptic transmission was triggered by one millisecond current injections using a concentric bipolar microelectrode (FHC; Model: CBAEC75) placed about 100-150 μm from the cell bodies of patched neurons. The extracellular stimuli were manipulated using an Isolated Pulse Stimulator (World Precision Instruments). Cells were held at −70V for all experiments. The evoked responses were measured by whole-cell recordings using a Multiclamp 700B amplifier (Molecular Devices). The whole-cell pipette solution contained 135 mM CsCl, 10 mM HEPES-CsOH (pH 7.25), 0.5 mM EGTA, 2 mM MgCl_2_, 0.4 mM NaCl-GTP, and 4 mM NaCl-ATP. The bath solution contained 140 mM, NaCl, 5 mM KCl, 2 mM CaCl_2_, 0.8 MgCl_2_, 10 mM HEPES-NaOH (pH 7.4), and 10 mM Glucose. For Aβ42 treatment peptide was prepared as described above. Neurons were treated with 500 nM Aβ42 24 hours prior to recording, and 500 nM Aβ42 was also added in bath solution during recording. EPSCs were distinguished by including 50 μM picrotoxin (Sigma) in the bath solution. The mEPSCs were sampled at 10 kHz in the presence of 1 μM tetrodotoxin (TTX, Sigma). The resistance of pipettes was 3-5 MΩ. The series resistance was adjusted to 8-10 MΩ once the whole-cell configuration was established.

**Antibodies:** Antibodies were obtained from Synaptic Systems, Goettingen, Germany (SYP, VAMP2, MAP2), Santa Cruz Biotechnology, Santa Cruz, California (synaptoporin, synaptogyrin1), and Pierce (synaptogyrin1), Covance Research Products, Inc. (Aβ42, 4G8 and 6E10) and Novus Biologicals (Aβ42, NB300-226).

**Human SYP expression and purification**. Full length 6xHis tagged human SYP was expressed using modified methods previously described for rat synaptophysin^38^. Isolated SF9 cells were solubilized with 1% Fos-choline 14 (FC14) and purified using a two-step chromatography employing a Ni-NTA column followed by a Sephadex S200 size exclusion column where human SYP eluted with an apparent molecule weight of ∼240kD corresponding to a hexamer. The samples were maintained in 0.15M sodium chloride, 0.03M sodium HEPES, pH 7.4, 0.009% FC14 at 4°C.

**Surface Plasmon Resonance:** Binding studies were performed on a Biacore 3000 or a BiOptix 404pi, with similar results. For the Biacore, a CM5 chip was used and for the BiOptix instrument, a CMV150 chip was employed. BioPure™ recombinant Aβ42, scrambled Aβ42, or Aβ42-1 (AmideBio, Boulder, CO) was dissolved to 0.1mM in 10mM NaOH, and diluted to 1uM in 10mM NaOAc pH 4.0 immediately prior to immobilization using EDC-NHS chemistry. Indicated concentrations of recombinant human SYP, containing a His(6)-tag was used as analyte. Bovine serum albumin (Sigma) was used as a control protein. Binding at a flow rate of 20μl/min was in 0.15M sodium chloride, 0.03M sodium HEPES, pH 7.4, 0.009% Fos-choline 14 for all samples.

**Immunoprecipitation:** Neurons were gently scraped from the dish in PBS and pelleted at 200 x g, then lysed in lysis buffer (320 mM sucrose, 10 mM HEPES pH 7.4, 1 mM EGTA, 100 μM EDTA, 0.1% (v/v) triton X-100) supplemented with protease inhibitor cocktail (Roche) rocking for 1 hr at 4 °C. Five μg of precipitating antibody was bound to PureProteome protein A magnetic beads (Millipore, Billerica, Massachusetts) in IP buffer (25 mM HEPES pH 7.4, 150 mM NaCl, 10 mM MgCl_2_, 1 mM EDTA, 1% glycerol, 0.5% NP-40) for 10 minutes and the beads were washed. Neuron lysates were applied in IP buffer with protease inhibitor cocktail (Roche) overnight at 4 °C. Beads were washed thrice in IP buffer and bound material was eluted at 95 °C in 2X SDS sample buffer.

**Densitometry:** Immunoblot results were quantified with ImageJ^36^. Levels of co-precipitated protein were normalized to levels of recovered bait protein and shown as a ratio over samples treated with scrambled Aβ42 (Fig. 2c), or normalized to MAP2 band and shown as ratio over WT protein levels (Fig. 2d).

**Alignments:** Human sequences of SYP and homologs were obtained from the Uniprot database (uniprot.org). Paralog tree was produced with the simple analysis tool from phylogeny.fr. ClustalW alignment was performed with gap penalties 12 open and 1 extension and aligned with PSI-BLAST at 3 iterations and an E-value cutoff of 0.01. This alignment was assigned a similarity score at each position by the PRALINE server using the BLOSUM62 matrix. The conservation scores were presented at each position as a moving average over a 3 residue window to smooth the plot. The query sequence (SYP) was then analyzed for hydropathy using the ExPASy ProtScale tool with the Kyte & Doolittle algorithm at a window size of 19 residues.

**Synaptosome preparation:** Whole brains were obtained from age-matched female B6 and *Syp*^*-/-*^ adults. Brains were homogenized 13 strokes on ice in 4 mL of sucrose buffer (10 mM HEPES pH 7.4, 320 mM sucrose, 2 mM EGTA, 2mM EDTA) with protease inhibitor cocktail (Roche) and homogenates were cleared at 4 °C at 1000g for 10 minutes. Synaptosomes were pelleted at 10,000g at 4 °C for 20 minutes, resuspended in buffer (25 mM HEPES pH 7.4, 150 mM NaCl, 10 mM MgCl_2_, 1 mM EDTA, 1% glycerol) and total protein was quantified with the Pierce 660 nM Protein Assay kit.

**Aβ42 affinity column:** AminoLink resin (Pierce, Rockford, Illinois) was functionalized according to manufacturer specifications with BioPure^(tm)^ recombinant Aβ42 or scrambled Aβ42 (AmideBio, Boulder, Colorado). Whole brain synaptosomes from B6 or *Syp*^*-/-*^ mice were applied to column overnight at 4 °C in IP buffer (25 mM HEPES pH 7.4, 150 mM NaCl, 10 mM MgCl_2_, 1 mM EDTA, 1% glycerol, 0.5% NP-40) with protease inhibitor cocktail (Roche). Beads were washed three times and bound material was eluted at 95°C in 2X SDS sample buffer.

**Purification of native SYP/VAMP2:** The native SYP/VAMP2 complex was purified from bovine brain tissue according to methods described previously^22^.

**Hippocampal slice preparation and electrophysiology:** Hippocampal slices (400 μm) were prepared from mice 2–4 months of age using a vibratome as described previously^39^. The slices were maintained at room temperature in a submersion chamber with artificial CSF (125 mM NaCl, 2.5 mM KCl, 2 mM CaCl_2_, 1 mM MgCl_2_, 1.25 mM NaH_2_PO_4_, 24 mM NaHCO_3_, and 15 mM glucose) bubbled with 95% O_2_/5% CO_2_. Slices were incubated for at least 2 h before removal for experiments. For electrophysiology experiments, slices were transferred to recording chambers (preheated to 32°C) where they were superfused with oxygenated ACSF. Monophasic, constant-current stimuli (100 μs) were delivered with a bipolar silver electrode placed in the stratum radiatum of area CA3, and the field EPSPs (fEPSPs) were recorded in the stratum radiatum of area CA1 with electrodes filled with ACSF (resistance, 2–4 MΩ). Baseline fEPSPs were monitored by delivering stimuli at 0.033 Hz. fEPSPs were acquired, and amplitudes and maximum initial slopes measured, using pClamp 10 (Molecular Devices). LTP was induced with a high-frequency stimulation (HFS) protocol consisting of two 1s long 100 Hz trains, separated by 60 s, delivered at 70–80% of the intensity that evoked spiked fEPSPs^40^. Incubation of hippocampal slices with Aβ42 was performed in either recording chambers or maintenance chambers as needed. The final concentrations of Aβ42 stock was prepared in DMSO and stored at −20°C for at least 24 h before use at a final concentration of 500 nm.

**Aβ42 ELISA:** Human Aβ42 was quantitated using a sandwich ELISA kits using the manufacturer’s protocol (Life Technologies). Briefly, whole brain extracts for each age group of both wild type (WT) and Tg-AD mice were analyzed and the level of Aβ42 was normalized to total protein as determined by BCA Assay.

**Pharmacological Seizure Susceptibility**. Mice of each genotype were assayed at 4-6 months of age. All mice were weighed the day of the experiment and were administered 25mg/kg kainic acid via i.p. from a freshly prepared 5mg/mL PBS solution. Mice were immediately placed in cylindrical observation chamber and monitored and scored in real time as well as constant video recording for 60 minutes. Both the observational and video recorded behavior were scored blind according to the modified Racine Scale.

## Acknowledgments

We thank our colleagues at the University of Colorado for helpful comments and criticisms, Rudolf Leube for the *Syp*^*-/-*^ mice, AmideBio for generous supply of BioPure^(tm)^ amyloid peptides, Domenico Galati for help with analysis of FM unloading data, Mary S. Rosendahl and BiOptix for access to BiOptix SPR instrumentation, and Brooke Hirsch and Robert S. Hodges for access to the Biacore 3000. Paula Villar and Maryam Amini for assistance the seizure experiments and Arieann DeFazio for help with the purification of SYP. This work was supported in part by an NIH EUREKA award (M.H.B.S.), an HHMI CIA award (M.H.B.S.), a Corden Pharma Fellowship (D.J.A.) and a NIH (T32 GM065103)/CU Molecular Biophysics Predoctoral Traineeship (D.J.A.).

### Author contributions

Experiments were conceived and designed by D.J.A. and M.H.B.S. Manuscript was written and edited by D.J.A. and M.H.B.S. Aβ42 binding column, VAMP2 coimmunoprecipitations, physin family analysis, primary neuronal culture and FM unloading experiments were performed and analyzed by D.J.A and M.H.B.S. Human SYP was expressed and purified by J.H.M. SPR experiments were performed and analyzed by S.P.E. and M.H.B.S. LTP experiments were performed by J.L. and analyzed by J.L., C.A.H. and M.H.B.S. Primary neuronal culture and electrophysiology was performed by C.S. and T.B. and analyzed by C.S. and M.H.B.S. Seizure susceptibility experiments were performed by L.H. and analyzed by L.H and M.H.B.S.

### Competing financial interests

The University of Colorado has filed a patent related to SYP as a potential target for the treatment of AD.

### Materials & correspondence

Correspondence and material requests should be addressed to MHBS.

### Extended data

**Extended Data Figure 1.**
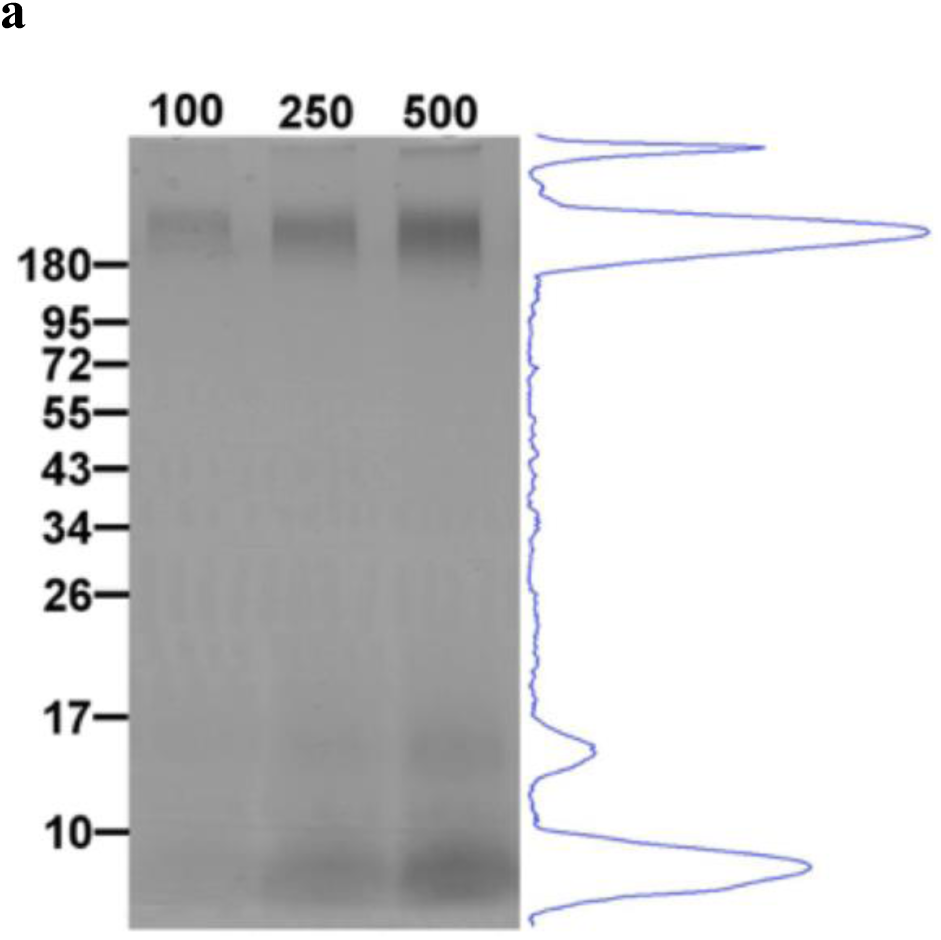
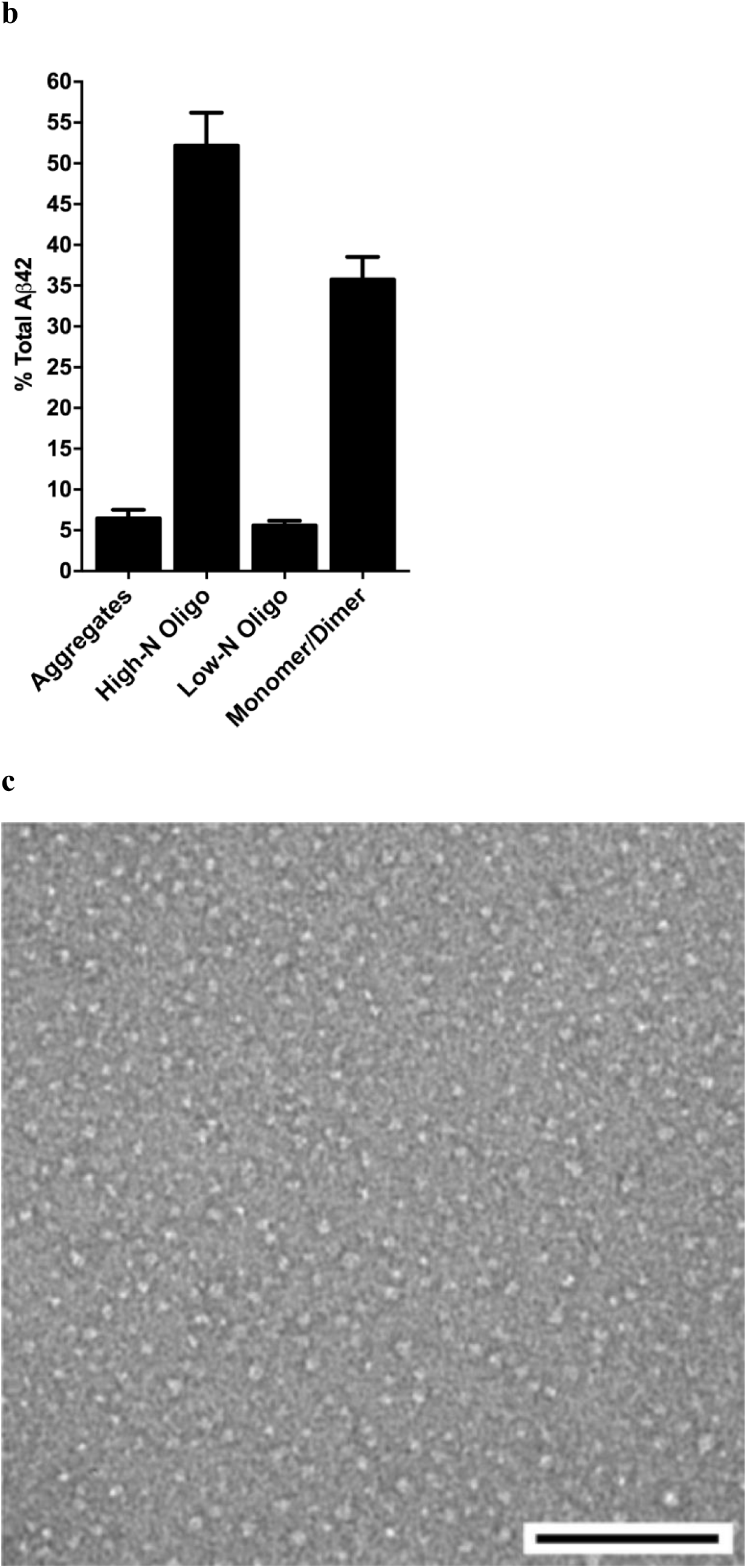
| Predominantly soluble oligomers and monomeric/dimeric Aβ42 used for neuron treatment and binding studies. Recombinant Aβ42 peptide was prepared fresh for each experiment from lyophilized film by first dissolving into 10 mM NaOH under sonication and then diluting in culture media. Prepared peptide was incubated for 10 minutes under experimental conditions prior to aggregation analysis. **a,** 100, 250 or 500 ng of peptide was assayed by native PAGE and silver staining to determine aggregation state. Molecular weight markers are in kDa and a line-scan plot of the 500 ng lane is shown at right. **b,** Line-scan data from multiple replicates (*n*=3) was used to estimate the relative levels of various peptide species present in the sample. **c,** Representative electron micrograph of negative stained peptide sample at 28K magnification confirms that the predominant forms observed under these conditions include small oligomers ranging from 5 to 8 nm in diameter (corresponding to the ∼200 kDa band in **a**), monomers, dimers and trimers (corresponding to the ∼5,10 and 15 kDa bands in **a**) which are too small to be observed under these conditions. Scale bar = 100 nm. Data are mean±s.e.m.

**Extended Data Figure 2.**
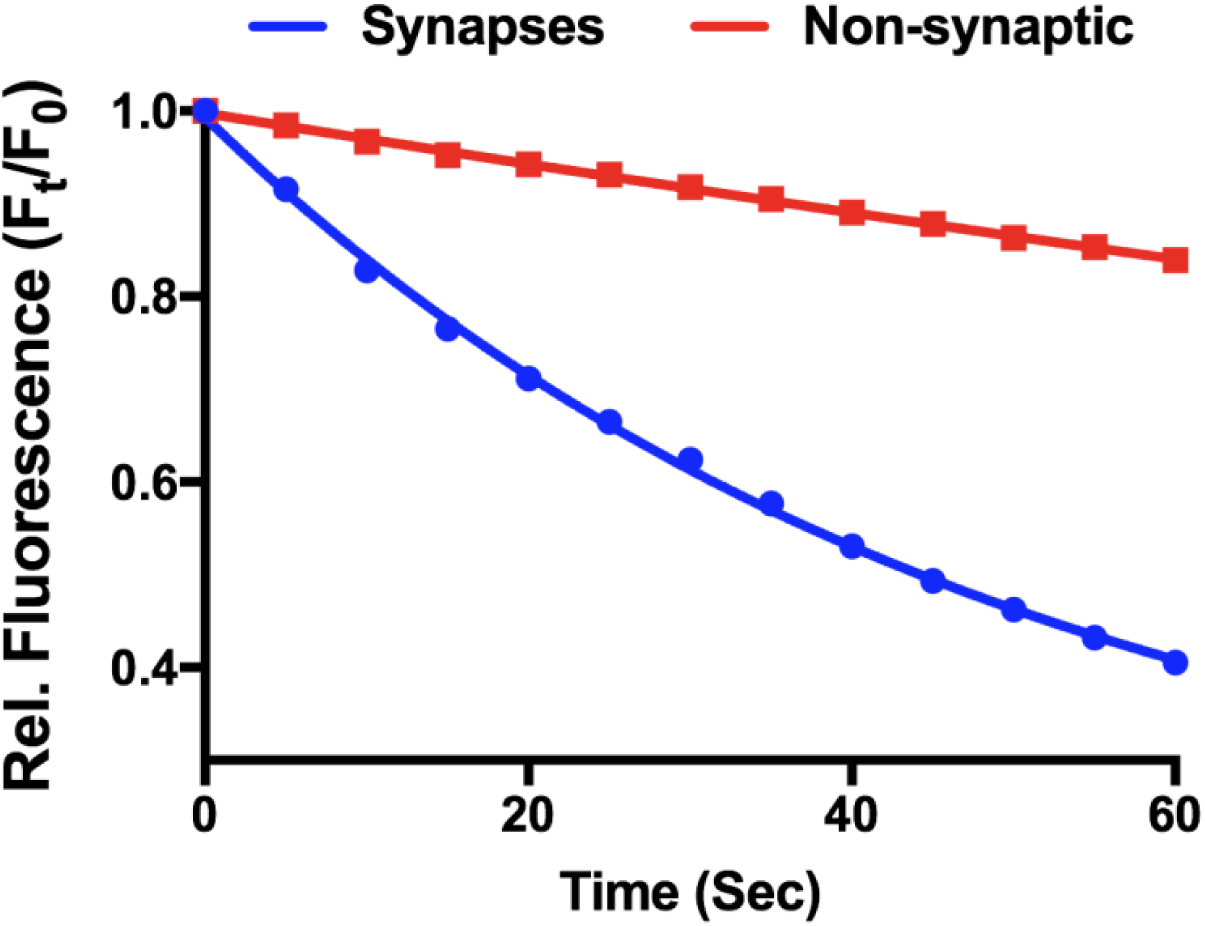
| FM Photobleaching rate is low relative to FM unloading rate. Relative fluorescence intensity of synaptic puncta and non-synaptic staining averaged across experiments shown in Fig. 1a during unloading. Non-synaptic background staining decays at photobleaching rate (τ=436s).

**Extended Data Figure 3.**
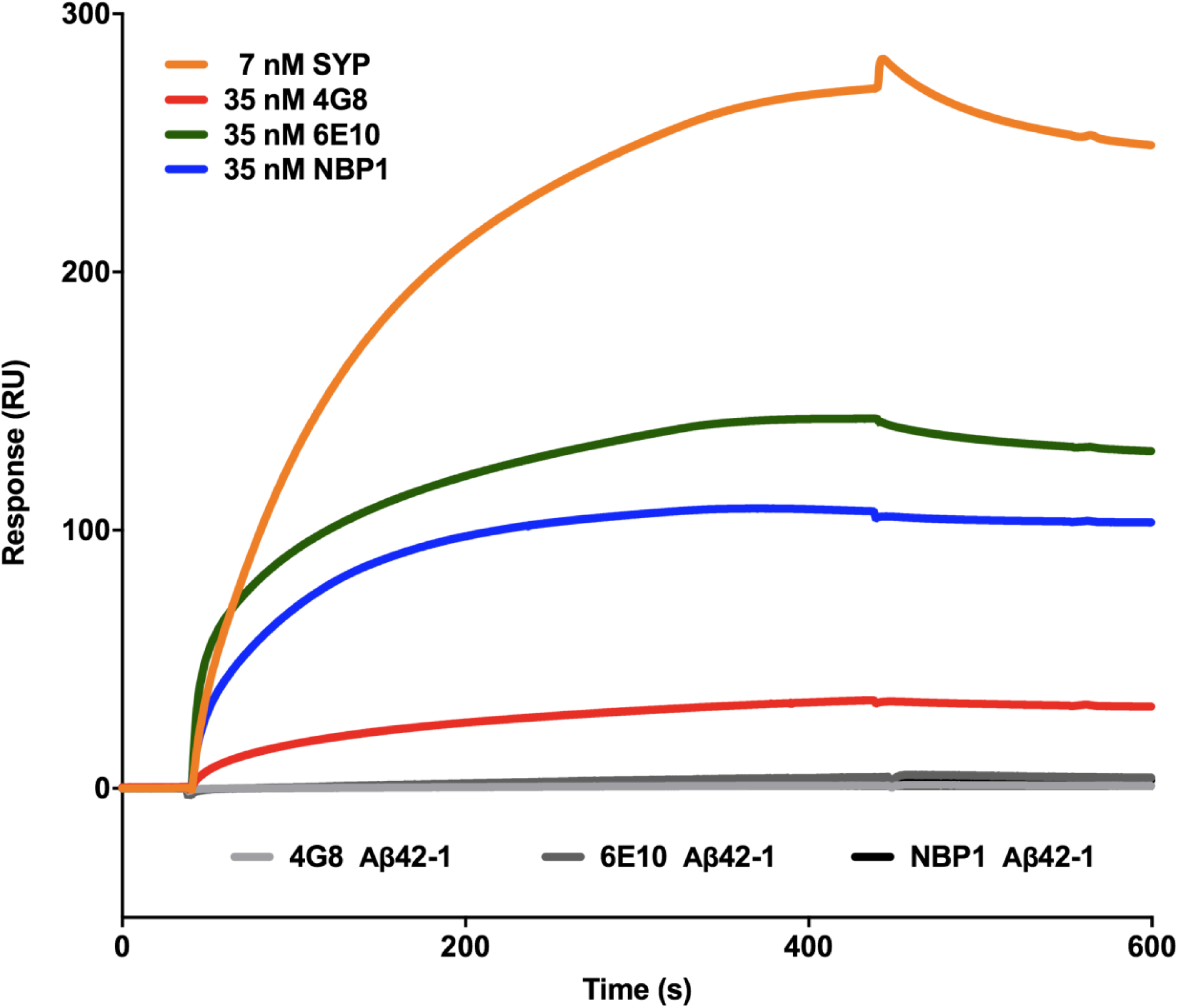
| Aβ42 on SPR chip is predominantly monomeric/dimeric and soluble oligomers. Antibodies with well characterized specificity for monomeric/dimeric (6E10), soluble oligomeric (NBP1) or protofibrillar (4G8) Aβ42 were used as analytes in an SPR experiment with Aβ42 prepared as described in Methods as ligand. Orange trace shows SYP binding for comparison. Grey traces show no binding to reverse Aβ42 peptide by any antibody confirming specificity.

**Extended Data Figure 4.**
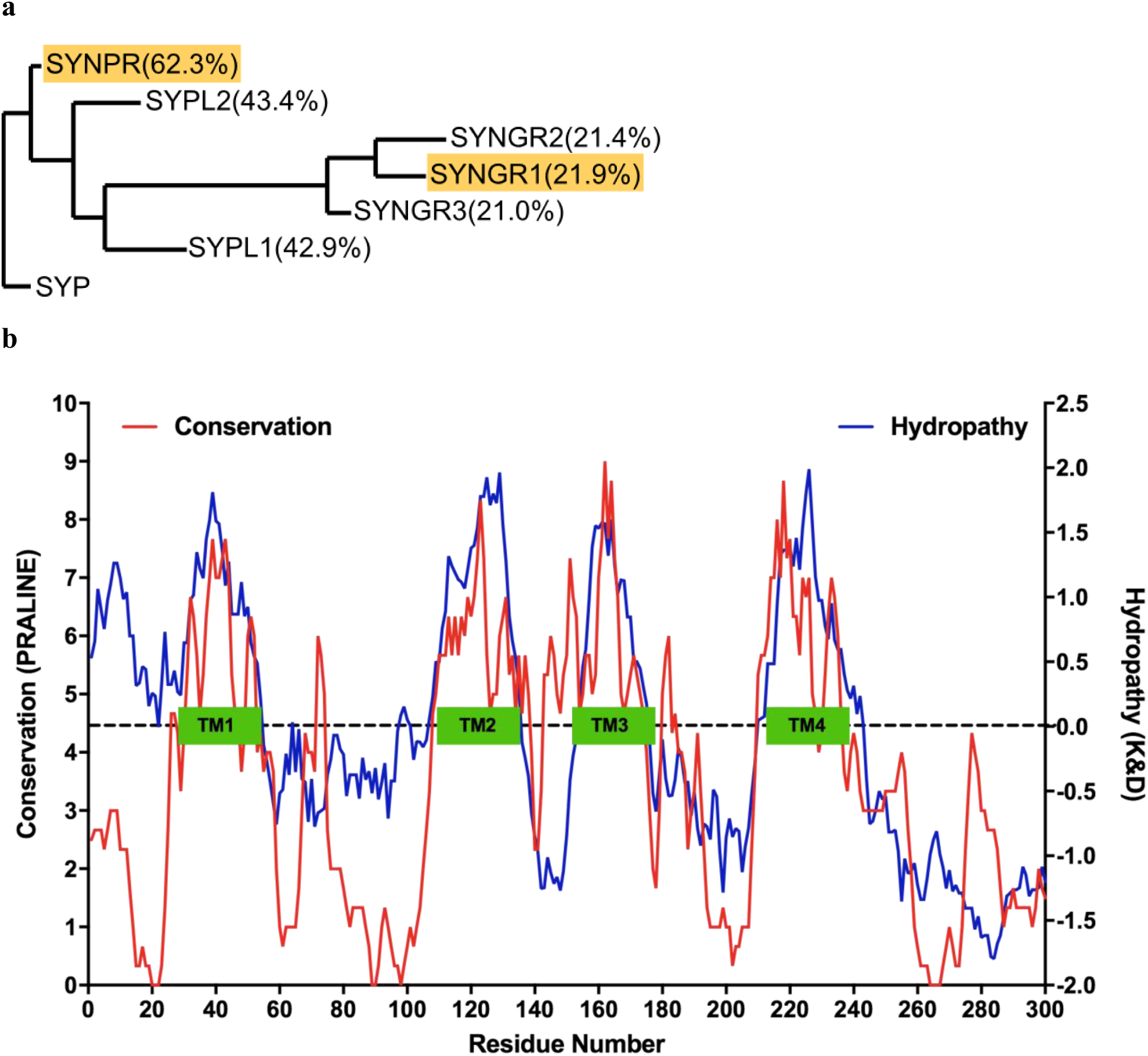
| SYP Paralogs most highly conserved in the transmembrane domains. **a,** PhyML tree for SYP and all 6 neuronal paralogs, % identity to SYP indicated in parentheses. Paralogs studied in Fig. 2 are highlighted in orange. **b,** Conservation score of physin family alignment (red) overlaid with SYP hydropathy score by the Kyte & Doolittle method (blue). TMDs indicated by green boxes.

**Extended Data Figure 5.**
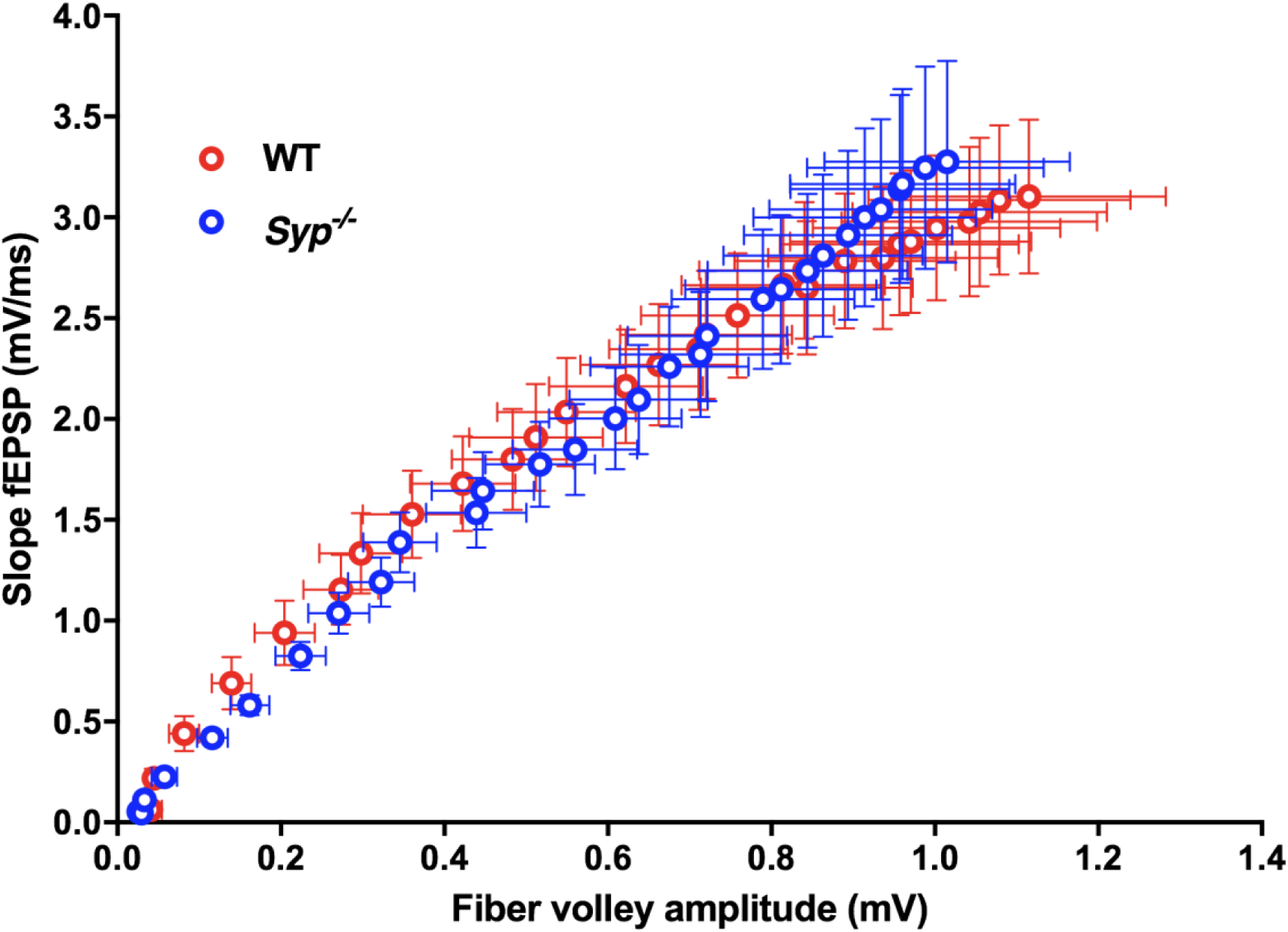
| Basal neurotransmission is normal in *Syp-/-* slices. Wild type and *Syp*^*-/-*^ mice show similar input/output ratios demonstrating no difference in the basal neurotransmission between WT and *Syp*^*-/-*^. Data are mean±s.e.m.

**Extended Data Figure 6.**
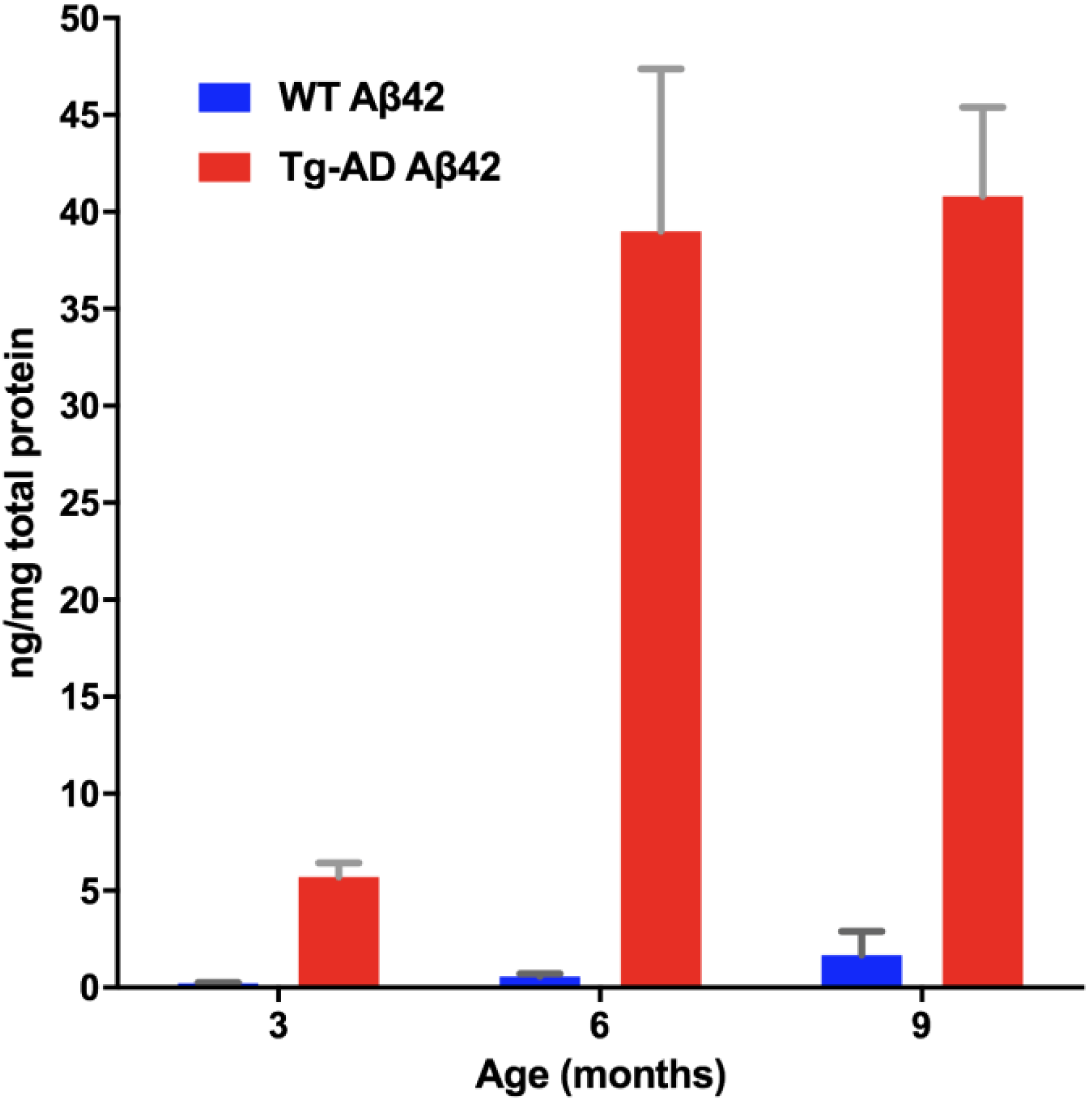
| TgAD mice accumulate Aβ42 in brains from 3 months on. Total brain levels of Aβ42 quantified by ELISA from WT and Tg-AD mice at 3, 6 and 9 months of age (*n*=4 for all groups). Data are mean±s.e.m.

